# Cross-species complementation reveals conserved functions for EARLY FLOWERING 3 between monocots and dicots

**DOI:** 10.1101/131185

**Authors:** He Huang, Malia A. Gehan, Sarah E. Huss, Sophie Alvarez, Cesar Lizarraga, Ellen L. Gruebbling, John Gierer, Michael J. Naldrett, Rebecca K. Bindbeutel, Bradley S. Evans, Todd C. Mockler, Dmitri A. Nusinow

## Abstract

Plant responses to the environment are shaped by external stimuli and internal signaling pathways. In both the model plant *Arabidopsis thaliana* and crop species, circadian clock factors have been identified as critical for growth, flowering and circadian rhythms. Outside of *A. thaliana,* however, little is known about the molecular function of clock genes. Therefore, we sought to compare the function of *Brachypodium distachyon* and *Seteria viridis* orthologs of *EARLY FLOWERING3,* a key clock gene in *A. thaliana.* To identify both cycling genes and putative ELF3 functional orthologs in *S. viridis*, a circadian RNA-seq dataset and online query tool (Diel Explorer) was generated as a community resource to explore expression profiles of *Setaria* genes under constant conditions after photo- or thermo-entrainment. The function of *ELF3* orthologs from *A. thaliana, B. distachyon,* and *S. viridis* were tested for complementation of an *elf3* mutation in *A. thaliana.* Despite comparably low sequence identity versus AtELF3 (less than 37%), both monocot orthologs were capable of rescuing hypocotyl elongation, flowering time and arrhythmic clock phenotypes. Molecular analysis using affinity purification and mass spectrometry to compare physical interactions also found that BdELF3 and SvELF3 could be integrated into similar complexes and networks as AtELF3, including forming a composite evening complex. Thus, we find that, despite 180 million years of separation, BdELF3 and SvELF3 can functionally complement loss of *ELF3* at the molecular and physiological level.

**One Sentence Summary:** Orthologs of a key circadian clock component ELF3 from grasses functionally complement the Arabidopsis counterpart at the molecular and physiological level, in spite of high sequence divergence.

## INTRODUCTION

Plants have developed sophisticated signaling networks to survive and thrive in diverse environments. Many plant responses are shaped, in part, by an internal timing mechanism known as the circadian clock, which allows for the coordination and anticipation of daily and seasonal variation in the environment (Greenham and McClung, 2015). Circadian clocks, which are endogenous oscillators with a period of approximately 24 hours, are critical for regulating the timing of physiology, development, and metabolism in all domains of life (Bell-Pedersen et al., 2005; Doherty and Kay, 2010; Edgar et al., 2012; Harmer, 2009; Wijnen and Young, 2006). In plants and blue-green algae, circadian clocks provide an experimentally observable adaptive advantage by synchronizing internal physiology with external environmental cues (Dodd et al., 2005; Ouyang et al., 1998; Woelfle et al., 2004). Currently, circadian oscillators are best understood in the reference plant *Arabidopsis thaliana*, in which dozens of clock or clock-associated components have been identified using genetic screens and non-invasive, luciferase-based oscillating reporters (Hsu and Harmer, 2014; Nagel and Kay, 2012). These morning-, afternoon-, and evening-phased clock oscillators form multiple interconnected transcription-translation feedback loops and compose a complex network (Hsu and Harmer, 2014; Pokhilko et al., 2012). The *A. thaliana* circadian clock regulates a significant portion of physiology, including photosynthesis, growth, disease resistance, starch metabolism, and phytohormone pathways (Covington et al., 2008; Graf et al., 2010; Harmer et al., 2000; Michael et al., 2008; Wang et al., 2011b), with up to 30% of gene expression under circadian control (Covington et al., 2008; Michael et al., 2008).

Within the Arabidopsis clock network, a tripartite protein complex called the evening complex (EC) is an essential component of the evening transcription loop (Huang and Nusinow, 2016b). The EC consists of three distinct proteins, EARLY FLOWERING 3 (ELF3), EARLY FLOWERING 4 (ELF4) and LUX ARRHYTHMO (LUX, also known as PHYTOCLOCK1), with transcript and protein levels peaking in the evening (Doyle et al., 2002; Hazen et al., 2005; Hicks et al., 2001; Nusinow et al., 2011; Onai and Ishiura, 2005). The EC plays a critical role in maintaining circadian rhythms, by repressing expression of key clock genes (Dixon et al., 2011; Helfer et al., 2011; Herrero et al., 2012; Kolmos et al., 2011; Mizuno et al., 2014). Loss-of-function mutation of any EC component in *A. thaliana* results in arrhythmicity of the circadian clock and causes excessive cellular elongation and early flowering regardless of environmental photoperiod (Doyle et al., 2002; Hazen et al., 2005; Hicks et al., 2001; Khanna et al., 2003; Kim et al., 2005; Nozue et al., 2007; Nusinow et al., 2011; Onai and Ishiura, 2005).

In *A. thaliana*, ELF3 directly interacts with ELF4 and LUX, functioning as a scaffold to bring ELF4 and LUX together (Herrero et al., 2012; Nusinow et al., 2011). Additional protein-protein interaction studies and tandem affinity purification coupled with mass spectrometry (AP-MS) have identified many ELF3-associating proteins and established ELF3 as a hub of a complex protein-protein interaction network, which consists of key components from the circadian clock pathway and light signaling pathways (Huang et al., 2016a; Huang and Nusinow, 2016b; Liu et al., 2001; Yu et al., 2008). In this network, ELF3 directly interacts with the major red light photoreceptor phytochrome B (phyB), and CONSTITUTIVE PHOTOMORPHOGENIC 1 (COP1), which is an E3 ubiquitin ligase required for proper regulation of photomorphogenesis and also interacts with phyB (Liu et al., 2001; Yu et al., 2008). The physical interaction among ELF3, phyB, and COP1, together with recruitment of direct interacting proteins to the network, provides biochemical evidence for cross-talk between circadian clock and light signaling pathways (Huang and Nusinow, 2016b). Although much work does translate from *A. thaliana* to other plant species, interaction between ELF3 and other proteins have yet to be tested in species outside *A. thaliana*. Whether evening complex-like protein assemblages or a similar ELF3-containing protein-protein interaction network exists in species outside *A. thaliana* is an interesting question to ask.

Identification and characterization of clock genes in diverse plant species has revealed that many clock components are broadly conserved (Filichkin et al., 2011; Khan et al., 2010; Lou et al., 2012; Song et al., 2010). Furthermore, comparative genomics analysis has found that circadian clock components are selectively retained after genome duplication events, suggestive of the importance of their role in maintaining fitness (Lou et al., 2012). Recently, mutant alleles of *ELF3* were identified associated with the selection of favorable photoperiodism phenotypes in several crops, such as pea, rice, soybean and barley (Faure et al., 2012; Lu et al., 2017; Matsubara et al., 2012; Saito et al., 2012; Weller et al., 2012; Zakhrabekova et al., 2012). These findings are consistent with the reported functions of *A. thaliana* ELF3 in regulating the photoperiodic control of growth and flowering (Hicks et al., 2001; Huang and Nusinow, 2016b; Nozue et al., 2007; Nusinow et al., 2011). However, opposed to the early flowering phenotype caused by *elf3* mutants in *A. thaliana,* pea, and barley (Faure et al., 2012; Hicks et al., 2001; Weller et al., 2012), loss of function mutation of the rice or soybean *ELF3* ortholog results in delayed flowering (Lu et al., 2017; Saito et al., 2012), suggesting ELF3-mediated regulation of flowering varies in different plant species. The molecular mechanisms underlying this difference have not been thoroughly elucidated.

*Brachypodium distachyon* is a C3 model grass closely related to wheat, barley, oats, and rice. *Setaria viridis*, is a C4 model grass closely related to maize, sorghum, sugarcane, and other bioenergy grasses. Both grasses are small, transformable, rapid-cycling plants with recently sequenced genomes, making them ideal model monocots for comparative analysis with Arabidopsis (Bennetzen et al., 2012; Brutnell et al., 2010). Computational analysis of *B. distachyon* has identified putative circadian clock orthologs (Higgins et al., 2010), including *BdELF3.* However, no such comparative analysis has been done systematically in *S. viridis* to identify putative orthologs of circadian clock genes. Therefore, we generated a RNA-seq time-course dataset to analyze the circadian transcriptome of *S. viridis* after either photo- or thermo-entrainment and developed an online gene-expression query tool (Diel Explorer) for the community. We found that the magnitude of circadian regulated genes in *S. viridis* is similar to other monocots after photo-entrainment, but much less after thermal entrainment. We further analyzed the functional conservation of SvELF3, together with previously reported BdELF3, by introducing both ELF3 orthologs into *A. thaliana elf3* mutant for physiological and biochemical characterization. We found that *B. distachyon* and *S. viridis* ELF3 can complement the hypocotyl elongation, flowering time and circadian arrhythmia phenotypes caused by the *elf3* mutation in *A. thaliana.* Furthermore, AP-MS analyses found that *B. distachyon* and *S. viridis* ELF3 were integrated into a similar protein-protein interaction network *in vivo* as their *A. thaliana* counterpart. Our data collectively demonstrated the functional conservation of ELF3 among *A. thaliana, B. distachyon* and *S. Viridis* is likely due to the association with same protein partners, providing insights of how ELF3 orthologs potentially function in grasses.

## RESULTS

### Identifying and cloning ELF3 orthologs from *B. distachyon* and *S. viridis*

ELF3 is a plant-specific nuclear protein with conserved roles in flowering and the circadian clock in multiple plant species (Faure et al., 2012; Herrero et al., 2012; Liu et al., 2001; Lu et al., 2017; Matsubara et al., 2012; Saito et al., 2012; Weller et al., 2012; Zakhrabekova et al., 2012). To identify ELF3 orthologs in monocots, we used the protein sequence of *Arabidopsis thaliana* ELF3 (AtELF3) to search the proteomes of two model monocots *Brachypodium distachyon* and *Setaria viridis* using BLAST (Altschul et al., 1990). Among the top hits, we identified a previously reported ELF3 homolog in *B. distachyon* (*Bradi2g14290.1, BdELF3*) (Calixto et al., 2015; Higgins et al., 2010) and two putative *ELF3* homologous genes *Sevir.5G206400.1* (referred as *SvELF3a*) and *Sevir.3G123200.1* (referred as *SvELF3b*) in *S. viridis.* We used Clustal Omega (http://www.ebi.ac.uk/Tools/msa/clustalo/) to conduct multiple sequence alignments of comparing protein sequences of ELF3 orthologs with that of AtELF3 (Sievers et al., 2011). *BdELF3*, *SvELF3a* and *SvELF3b* encode proteins with similar identity compared to AtELF3 (34.7–36.8%) (**Supplemental Figure S1**). When compared to BdELF3, SvELF3b was 74.3% identical while SvELF3a was 57.4% identical (**Supplemental Figure S1**). Therefore, to maximize the diversity of ELF3 sequences used in this study, we cloned full length cDNAs encoding BdELF3 and SvELF3a.

### Diel Explorer of *S. viridis* circadian data

In Arabidopsis, ELF3 cycles under diel and circadian conditions (constant condition after entrainment) with a peak phase in the evening (Covington et al., 2001; Hicks et al., 2001; Nusinow et al., 2011). We queried an available diurnal time course expression dataset for *B. distachyon* from the DIURNAL website, and found that *BdELF3* expression cycles under diel conditions (LDHH, 12 h light / 12 h dark cycles with constant temperature), but not under circadian conditions in available data (LDHC-F or LDHH-F, **Supplemental Figure S2**) (Filichkin et al., 2011; Mockler et al., 2007). Also different from *AtELF3,* transcript levels of *BdELF3* accumulate at dawn rather than peak in the evening (**Supplemental Figure S2**) when grown under diel conditions, suggesting different regulations on *ELF3* expression between monocot and dicot plants. Neither diel nor circadian expression data for *S. viridis* was available. Therefore, we generated RNA-seq time-course data to examine *SvELF3* expression as well as the circadian expression of other clock orthologs after both photocycle and thermocycle entrainment. In addition, we developed the Diel Explorer tool(http://shiny.bioinformatics.danforthcenter.org/diel-explorer/) to query and visualize *S. viridis* circadian-regulated gene expression (**Supplemental Figure S3**). 48,594 *S. viridis* transcripts are represented in the two datasets entrained under either photocycles (LDHH-F) or thermocycles (LLHC-F). With Diel Explorer users can manually enter a list of transcript identifiers, gene ontology (GO) terms, or gene orthologs, plot gene expression, and download data. Alternatively, users can upload files of transcript identifiers or gene orthologs, and/or filter the datasets by entrainment, phase, or significance cut-offs. Data and graphs can be downloaded directly using Diel Explorer. The tool serves as a community resource that can be expanded to include other circadian or diurnal data in the future. The underlying code is available on Github (https://github.com/danforthcenter/diel-explorer).

Under photoperiod entrainment (LDHH-F), 5,585 of the 48,594 *S. viridis* transcripts are circadian regulated (Bonferroni-adjusted P-Value < 0.001). This proportion of photoperiod-entrained circadian genes (~11.5%) is similar to maize (10.8%), rice (12.6%), and poplar (11.2%) data sets, but much smaller than the approximately 30% reported for *A. thaliana* (Covington et al., 2008; Filichkin et al., 2011; Khan et al., 2010). Under thermocycle entrainment (LLHC-F), 582 of the 48,594 *S. viridis* transcripts are circadian regulated. Therefore, only ~1.2% of *S. viridis* transcripts are circadian cycling under thermocycle entrainment. The ~10-fold reduction in circadian cycling genes between photocycle and thermocycle entrainment (**Supplemental Figure S4**) is interesting considering that there was less than 1% difference in the number of genes with a circadian period between photocycle and thermocycle entrainment in C3 monocot rice (*Oryza japonica*) (Filichkin et al., 2011). The reduction in cycling genes between the two entrainment conditions in *S. viridis* compared to *O. japonica* is an indication that circadian regulation could vary greatly among monocots. Also, the difference in number of cycling genes between monocots and dicots may represent a significant reduction of the role of the circadian clock between these lineages.

In addition to the overall reduction in circadian genes, the phase with the most number of cycling genes was ZT18 after light entrainment (LDHH-F; **Supplemental Figure S4**), but ZT12 with temperature entrainment (LLHC-F; **Supplemental Figure S4**), which is consistent with previous studies that have found significant differences in temperature and light entrainment of the circadian clock (Boikoglou et al., 2011; Michael et al., 2008; Michael et al., 2003). There are 269 genes that are considered circadian-regulated and are cycling under both LDHH-F and LLHC-F conditions (Bonferroni Adjusted P-Value < 0.001). The list of 269 genes that overlap between photocycle and thermocycle entrainment includes best matches for Arabidopsis core clock components *TIMING OF CAB EXPRESSION 1* (*TOC1*, *AT5G61380.1*; *Sevir.1G241000.1*), *LATE ELONGATED HYPOCOTYL* (*LHY*, *AT1G01060*; *Sevir.6G053100.1*), and *CCA1*-like gene *REVEILLE1* (*RVE1*, *AT5G17300.1*; *Sevir.1G280700.1*). However, putative *S. viridis* orthologs of *TOC1*, *LHY*, and *RVE1* all have different circadian phases under LDHH entrainment compared to LLHC entrainment (**Figure 1**). In fact, the majority (233/269) of overlapping circadian genes in *S. viridis* have a distinct circadian phase under thermocyle compared to photocyle entrainment (**Supplemental Figure S4**). We also found that putative orthologs of *PSEUDO-RESPONSE REGULATOR 7* (*PRR7*, *AT5G02810*; *Sevir.2G456400.1*; related to *OsPRR73* (Murakami et al., 2003)) and *LUX ARRHYTHMO* (*LUX*, *AT3G46640*; *Sevir.5G474200.1*) cycle significantly under LDHH-F but not LLHC-F conditions (**Figure 1**). Neither *SvELF3a* nor *SvELF3b* cycle under circadian conditions after photo- or thermo-entrainment (**Figure 1**), similar to ELF3 orthologs in *B. distachyon* (**Supplemental Figure S2**) (Mockler et al., 2007) and *O. sativa* (Filichkin et al., 2011). This is different from *AtELF3*, which continues to cycle under constant condition after either photo- or thermos-entrainment (**Supplemental Figure S5**) (Mockler et al., 2007). The difference in expression of these putative orthologs between *A. thaliana* and monocots *S. viridis*, *B. distachyon*, and *O. sativa*, suggest that the architecture of the circadian clock may have significant differences in response to environmental cues in these two species.

**Figure 1.**
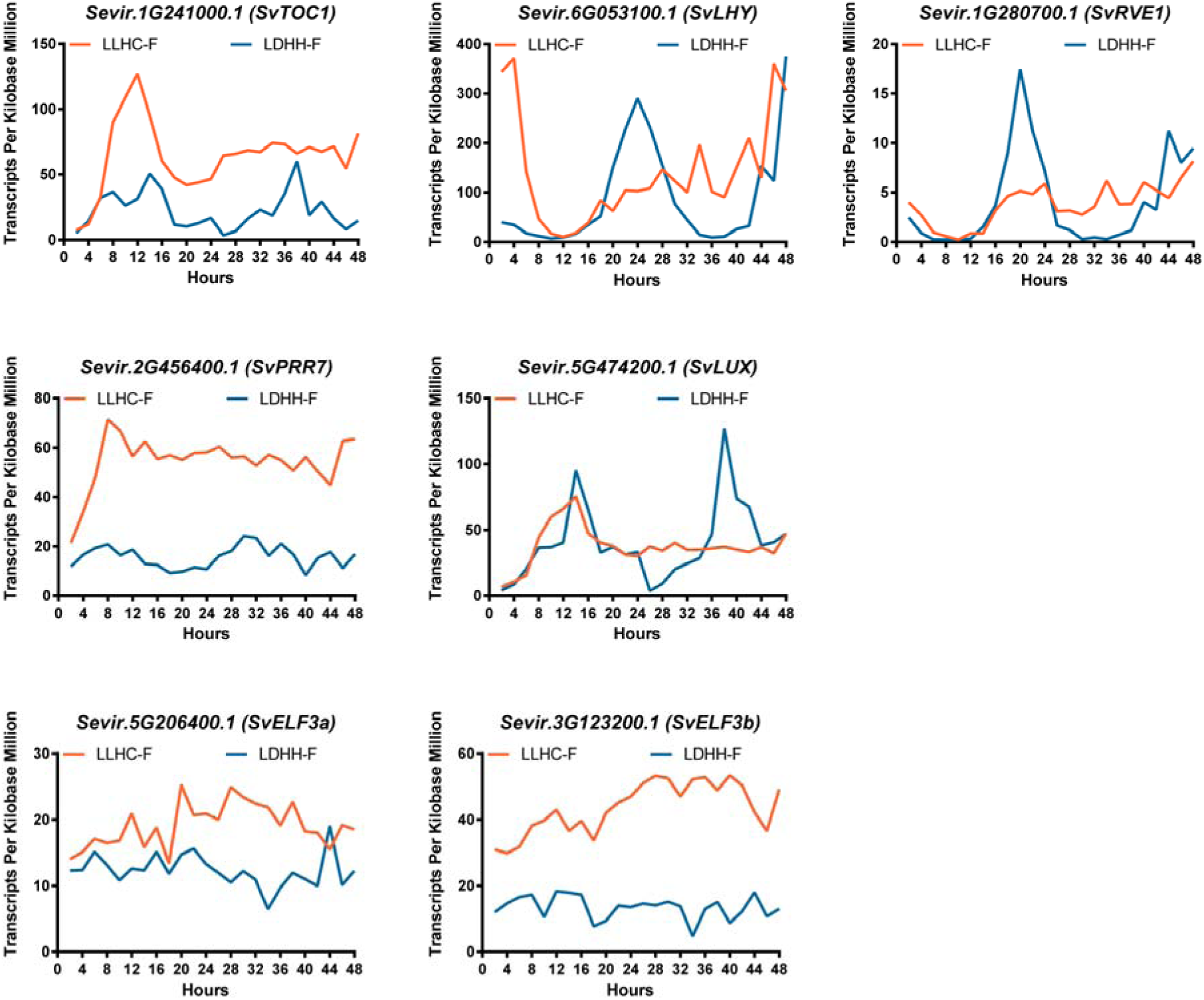
Circadian expression profiles of putative *S. viridis* clock components from Diel Explorer using time-course RNA-seq data. *S. viridis* plants were entrained by either photocycle (LDHH) or thermocycle (LLHC), followed by being sampled every 2 hours for 48 hours under constant temperature and light conditions (Free-Running; F) to generate time-course RNA-seq data. Mean values of Transcripts per Kilobase Million (TPM) from two experimental replicates for each timepoints per gene were plotted.

### BdELF3 and SvELF3 rescue growth and flowering defects in *Arabidopsis elf3* mutant

Although the circadian expression pattern of *B. distachyon ELF3* and *S. viridis ELF3* is different from that of *A. thaliana ELF3*, it is still possible that the ELF3 orthologs have conserved biological functions. To test this, we sought to determine if BdELF3 or SvELF3a could complement the major phenotypic defects of the *elf3* mutant in *A. thaliana*, namely hypocotyl elongation, time to flowering, or circadian rhythmicity. To this end, we constitutively expressed *BdELF3*, *SvELF3a* (hereafter referred as *SvELF3*) and *AtELF3* cDNAs by the *35S Cauliflower mosaic virus* promoter in the *A. thaliana elf3-2* mutant expressing a *LUCIFERASE* reporter driven by the promoter of *CIRCADIAN CLOCK ASSOCIATED 1* (*CCA1*) (*elf3-2* [*CCA1:LUC*]) (Pruneda-Paz et al., 2009). All three ELF3 coding sequences were fused to a C-terminal His_6_-3xFlag affinity tag (HFC), which enables detection by western blotting and identification of protein-protein interaction by affinity purification and mass spectrometry (AP-MS) (Huang et al., 2016a). After transforming these constructs, we identified and selected two biologically independent transgenic lines with a single insertion of each At/Bd/SvELF3-HFC construct. Western blot analysis using FLAG antibodies detected the expression of all ELF3-HFC fusion proteins (**Supplemental Figure S6**).

Next, we asked if expressing At/Bd/SvELF3-HFC fusion proteins could rescue the mutation defects caused by *elf3-2*. When plants are grown under light/dark cycles (12 hour light: 12 hour dark), *elf3-2* mutant plants elongate their hypocotyls much more than wild type plants (4.75±0.48 mm vs. 1.95±0.27 mm, respectively. ± = standard deviation) (**Figure 2**). The long hypocotyl defect in *elf3-2* was effectively suppressed by expressing either *AtELF3*, or *ELF3* orthologs (*BdELF3 or SvELF3a*) (**Figure 2**). These data show that the monocot *ELF3* orthologs function similarly to *A. thaliana ELF3* in the regulation of hypocotyl elongation in seedlings.

**Figure 2.**
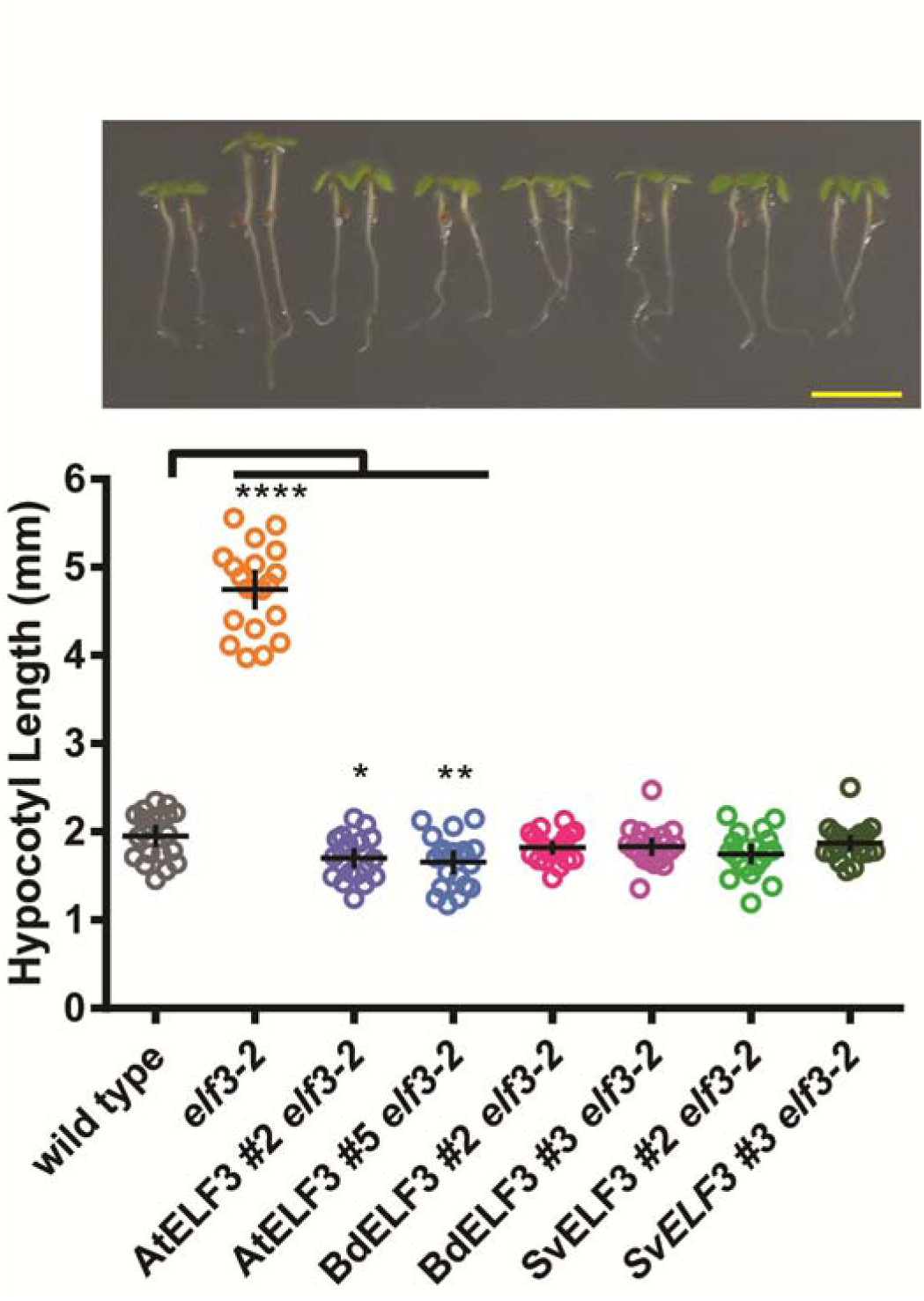
ELF3 orthologs suppress hypocotyl elongation defects in *elf3-2.* The hypocotyls of 20 seedlings of wild type, *elf3-2* mutant, AtELF3 *elf3-2*, BdELF3 *elf3-2*, and SvELF3 *elf3-2* (two independent transgenic lines for each ELF3 ortholog) were measured at 4 days after germination under 12-hour light :12-hour dark growth conditions at 22 °C. Upper panel shows representative seedlings of each genotype, with scale bar equal to 5 mm. Mean and 95% confidence intervals are plotted as crosshairs. This experiment was repeated three times with similar results. ANOVA analysis with Bonferroni correction was used to generate adjusted P values, * < 0.05, ** < 0.01, **** < 0.0001.

In addition to regulating phenotypes in seedlings, ELF3 also functions in adult plants to suppress the floral transition. Loss-of-function in Arabidopsis *ELF3* results in an early flowering phenotype regardless of day-length (Hicks et al., 2001; Liu et al., 2001; Zagotta et al., 1992). To determine how monocot ELF3 orthologs compared to *A. thaliana* ELF3 in flowering time regulation, we compared flowering responses under long day conditions among wild type, *elf3-2*, and *elf3-2* transgenic lines expressing *AtELF3, BdELF3,* or *SvELF3* (At/Bd/SvELF3-HFC). Constitutive over-expression of *AtELF3* led to a delay in flowering in long days (**Figure 3**) as previously observed (Liu et al., 2001). Similarly, constitutive expression of *BdELF3* or *SvELF3* caused plants to flower significantly later than the *elf3* mutants. These data show that all *ELF3* orthologs can function to repress the rapid transition to flowering of the *elf3* mutation when constitutively expressed in adult plants.

**Figure 3.**
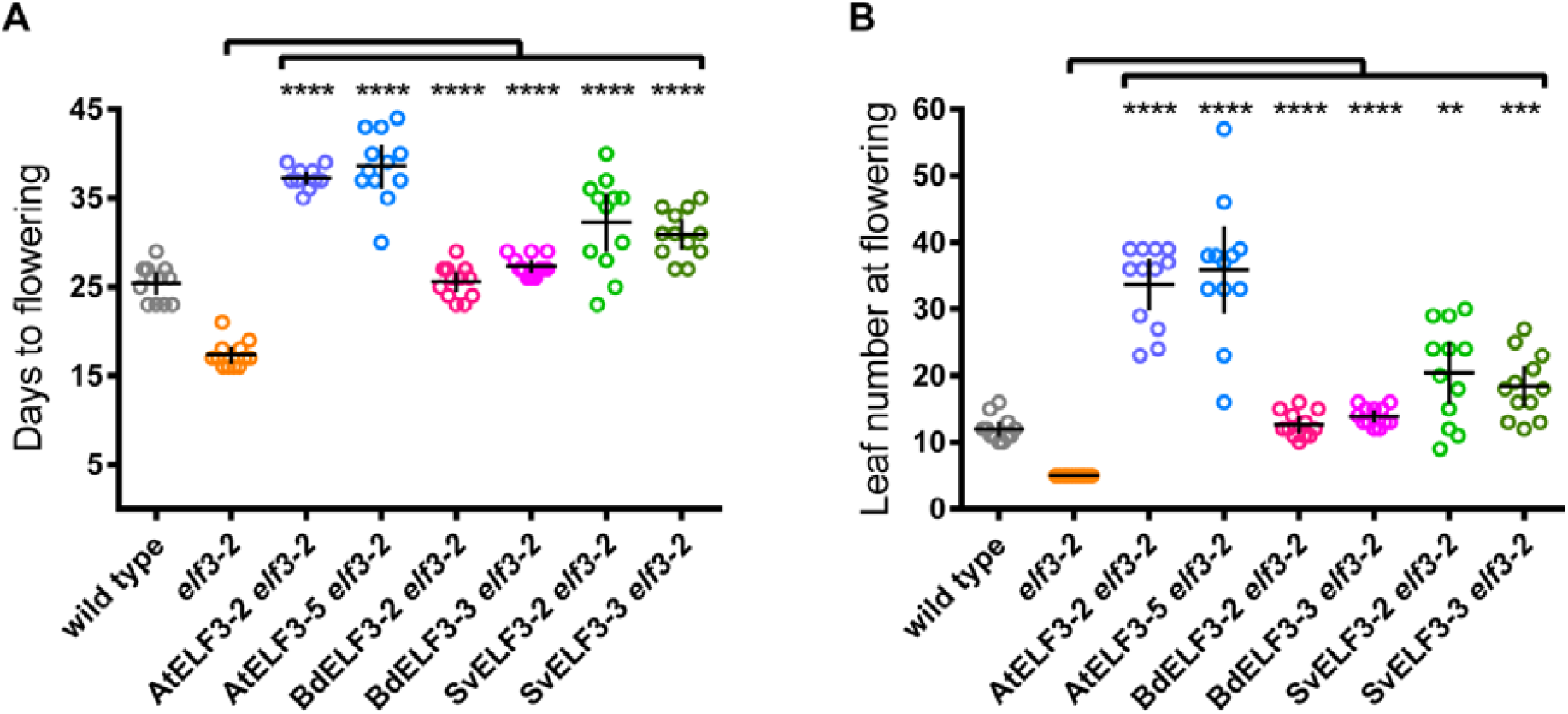
ELF3 orthologs suppress time to flowering of *elf3-2.* 12 wild type, *elf3-2* mutant, AtELF3 *elf3-2*, BdELF3 *elf3-2*, and SvELF3 *elf3-2* seedlings from two independent transformations were measured for days **(A)** and number of rosette leaves **(B)** at flowering (1 cm inflorescence). Mean and 95% confidence intervals are plotted as crosshairs. This experiment was repeated twice with similar results. ANOVA analysis with Bonferroni correction was used to generate adjusted P values, ** < 0.01, *** < 0.001, **** < 0.0001, of measurements when compared to the *elf3-2* mutant line.

### BdELF3 and SvELF3 restore the circadian rhythmicity in *Arabidopsis elf3* mutant

ELF3 is a key component of the *A. thaliana* circadian clock and is critical for maintaining the periodicity and amplitude of rhythms as shown using the *CCA1* promoter driven luciferase reporter (CCA:LUC) (Covington et al., 2001; Hicks et al., 1996; Nusinow et al., 2011). To determine if *BdELF3* or *SvELF3*a could rescue the arrhythmic phenotype of the *elf3* mutation, we analyzed the rhythms of the *CCA1::LUC* reporter under constant light conditions after diel entrainment (12 hours light: 12 hours dark at constant 22 °C). Relative amplitude error (RAE) analysis found that 100% of wild type and nearly all of the three *elf3-2* transgenic lines expressing *AtELF3, BdELF3* (*BdELF3 #3* was 87.5% rhythmic), and *SvELF3* were rhythmic, while only 37.5% of the *elf3-2* lines had rhythms (RAE < 0.5) (**Supplemental Figure S7**). Comparison of average period length found that the *AtELF3* expressing lines completely rescued the period and amplitude defects in the *elf3* mutant (**Figure 4A**, and **4D**). The *SvELF3* and *BdELF3* lines also rescued the amplitude defect (**Figure 4B** and Figure 4C), but their period was longer than wild type (compare 23.21 ± 0.59 hours for wild type to *BdELF3 #2*= 27.68 ± 0.86 hours, *BdELF3 #3*= 26.39 ± 1.01 hours, *SvELF3 #2*= 24.47 ± 0.50 hours, *SvELF3 #3*= 24.32 ± 0.34 hours, and *elf3-2=* 31.27 ± 1.93 hours, ± = standard deviation, **Figure 4D**). In summary, these data show that expression of any of the ELF3 orthologs is sufficient to recover the amplitude and restore rhythms of the *CCA1::LUC* reporter.

**Figure 4.**
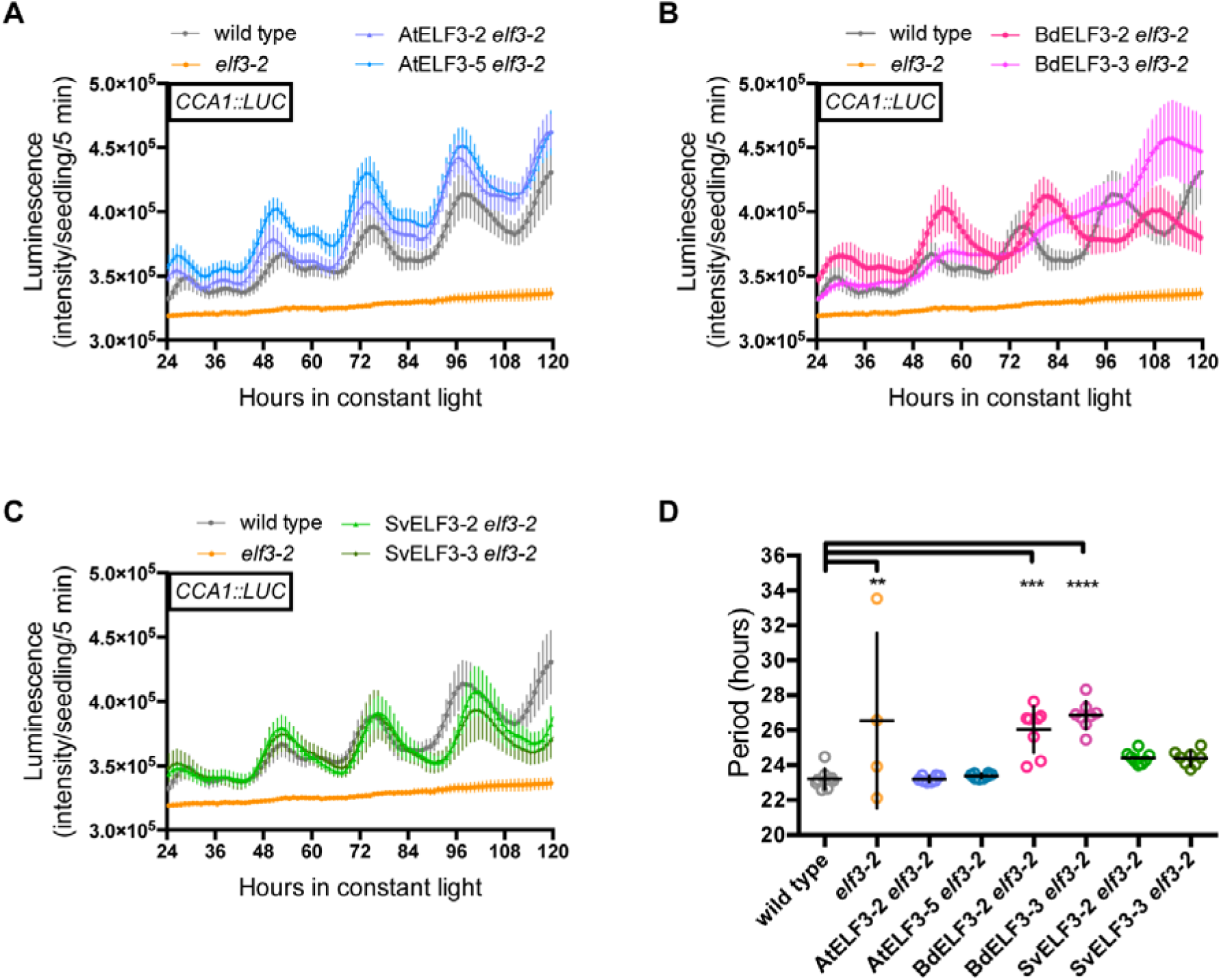
ELF3 orthologs can recover *CCA1::LUC* rhythms and amplitude in *elf3-2* mutants. 8 seedlings of wild type, *elf3-2* mutant, AtELF3 *elf3-2* **(A)**, BdELF3 *elf3-2* **(B)**, and SvELF3 *elf3-2* **(C)** from two independent transformations were imaged for bioluminescence under constant light after entrainment in 12-hour light :12-hour dark growth conditions at 22 °C. Each plot shows average bioluminescence of all seedlings along with 95% confidence interval (error bars). This experiment was repeated four times with similar results. Note that wild type and *elf3-2* mutant data was plotted on all graphs for comparison. **(D)** Periods of seedlings. Only periods with a Relative Amplitude Error below 0.5 (see Supplemental Figure S7) were plotted. Mean and 95% confidence intervals are plotted as crosshairs. ANOVA analysis with Bonferroni correction was used to generate adjusted P values, * < 0.05, ** < 0.01, *** < 0.001, **** < 0.0001, of measurements when compared to the wild type.

### BdELF3 and SvELF3 are integrated into a similar protein-protein interaction network in *A. thaliana*

Despite relatively low sequence conservation at the protein level, the *ELF3* orthologs can complement a wide array of *elf3* phenotypes (**Figures 2** to **4**). As ELF3 functions within the evening complex (EC) in Arabidopsis, which also contains the transcription factor LUX and the DUF-1313 domain containing protein ELF4 (Herrero et al., 2012; Nusinow et al., 2011), we reasoned that the monocot *ELF3* orthologs may also be able to bind to these proteins when expressed in *A. thaliana*. To determine if a composite EC could be formed, we tested if BdELF3 or SvELF3a could directly interact with AtLUX or AtELF4 in a yeast two-hybrid assay. Similar to AtELF3 (Nusinow et al., 2011), both BdELF3 and SvELF3a directly interact with both AtELF4 and the C-terminal portion of AtLUX (**Figure 5**). We cannot conclude that whether monocot ELF3 orthologs are also able to interact with the N-terminal AtLUX, since this fragment auto-activated the reporter gene in the yeast two-hybrid assay (**Figure 5**).

**Figure 5.**
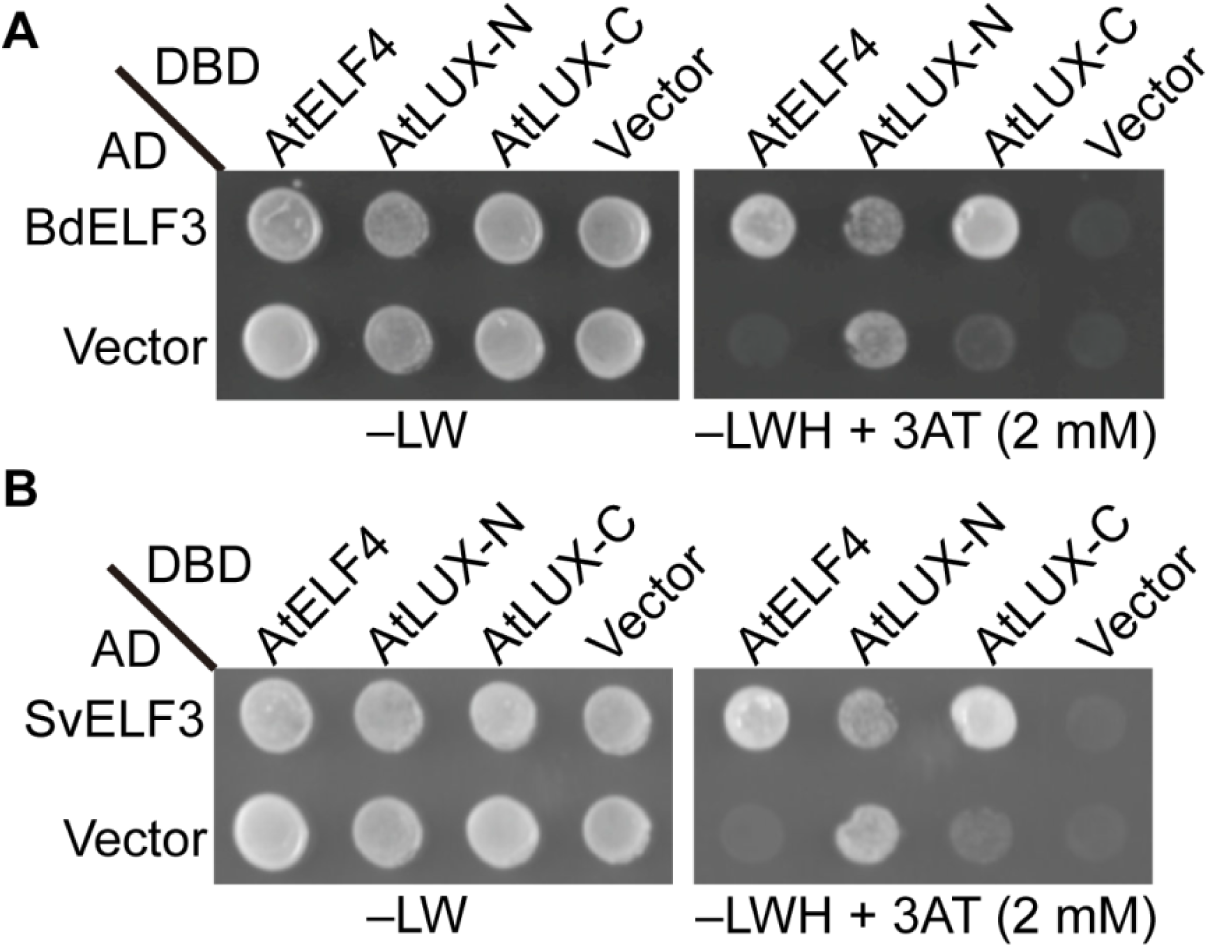
Both BdELF3 and SvELF3 can directly bind to AtELF4 and AtLUX. Yeast two-hybrid analysis of testing if either BdELF3 **(A)** or SvELF3 **(B)** can directly interact with either AtELF4, the N-terminal half of AtLUX (AtLUX-N, a.a. 1-143) or the C-terminal half of AtLUX (AtLUX-C, a.a. 144-324). –LW tests for the presence of both bait (DBD) and pray (AD) vectors, while the –LWH + 3AT tests for interaction. Vector alone serves as interaction control. This experiment was repeated twice with similar results.

ELF3 functions not only as the scaffold of the EC, but also as a hub protein in a protein-protein interaction network containing multiple key regulators in both the circadian clock and light signaling pathways (Huang et al., 2016a; Huang and Nusinow, 2016b). We hypothesize that BdELF3 and SvELF3 could rescue many of the defects of the *elf3* mutant because both monocot versions were integrated into the same protein-protein interaction network. To test this hypothesis, we used affinity purification and mass spectrometry (AP-MS) to identify the proteins that co-precipitate with monocot ELF3s when expressed in *A. thaliana*. AP-MS on two biological replicates for each sample with the above-mentioned independent insertion lines were included for each ELF3 ortholog. For comparison, the same AP-MS experiment was done with one of the 35S promoter-driven AtELF3-HFC transgenic lines (AtELF3-2). To detect specific co-precipitating proteins, we manually removed commonly identified contaminant proteins from plant affinity purifications and mass spectrometry experiments (Van Leene et al., 2015), and proteins identified from a control transgenic line expressing GFP-His_6_-3xFlag described previously (Huang et al., 2016a) (**Table 1**, the full list of identified proteins can be found in **Supplemental Table S2**).

**Table 1.** Proteins co-purified with ELF3 orthologs from AP-MS

| AGI number | Protein Name | Molecular Weight | Exclusive Unique Peptide Count/Percent Coverage <sup>a</sup> |  |  |  |  |  |  |  |  |  |
| --- | --- | --- | --- | --- | --- | --- | --- | --- | --- | --- | --- | --- |
|  |  |  | AtELF3 |  | SvELF3 #2 |  | SvELF3 #3 |  | BdELF3 #2 |  | BdELF3 #3 |  |
|  |  |  | rep1 | rep2 | rep1 | rep2 | rep1 | rep2 | rep1 | rep2 | rep1 | rep2 |
| n/a | AtELF3-HFC | 84 kDa | 20/36% | 19/31% | — | — | — | — | — | — | — | — |
| n/a | SvELF3-HFC | 87 kDa | — | — | 19/40% | 29/48% | 22/42% | 33/54% | — | — | — | — |
| n/a | BdELF3-HFC | 86 kDa | — | — | — | — | — | — | 25/42% | 26/43% | 21/35% | 19/33% |
| AT2G18790 | phyB | 129 kDa | 23/37% | 24/37% | 31/54% | 28/42% | 32/50% | 30/45% | 22/34% | 22/33% | 24/40% | 24/37% |
| AT5G35840 | phyC | 124 kDa | 22/29% | 23/29% | 27/37% | 32/39% | 27/37% | 33/42% | 20/24% | 18/22% | 19/23% | 22/27% |
| AT5G43630 | TZP | 91 kDa | 14/21% | 12/17% | 13/23% | 20/33% | 13/22% | 24/36% | 12/18% | 13/19% | 14/20% | 14/22% |
| AT4G18130 | phyE | 123 kDa | 11/16% | 19/27% | 12/18% | 18/23% | 11/19% | 17/24% | 6/7% | 10/14% | 14/19% | 13/18% |
| AT2G16365.2 | PCH1<sup>b</sup> | 51 kDa | 9/25% | 9/25% | 9/30% | 14/43% | 11/36% | 16/49% | 9/26% | 9/28% | 11/32% | 11/32% |
| AT2G46340 | SPA1 | 115 kDa | 8/14% | 7/9% | 2/2% | 4/8% | 2/3% | 4/9% | — | — | 6/8% | 5/6% |
| AT3G42170 <sup>c</sup> | DAYSLEEPER | 79 kDa | 8/19% | 5/11% | — | 7/16% | — | 12/27% | 3/7% | 4/8% | 2/4% | — |
| AT2G32950 | COP1 | 76 kDa | 8/16% | 9/17% | 6/17% | 4/8% | 3/6% | 5/12% | 1/2% | 2/4% | 6/11% | 4/8% |
| AT3G22380 | TIC | 165 kDa | 5/5% | 5/5% | 4/4% | 12/12% | 3/3% | 15/15% | 4/4% | 3/3% | 1/1% | 1/1% |
| AT4G11110 | SPA2 | 115 kDa | 5/11% | 6/12% | — | — | — | — | — | — | — | — |
| AT2G40080 | ELF4 | 12 kDa | 4/50% | 4/50% | 3/42% | 5/68% | 4/60% | 4/60% | 4/50% | 4/50% | 5/68% | 5/68% |
| AT1G09340 <sup>c</sup> | CRB | 43 kDa | 4/16% | 3/12% | 1/4% | 1/4% | 1/4% | 4/18% | 1/4% | 1/4% | — | — |
| AT3G13670 | MLK4 | 79 kDa | 3/10% | 3/13% | — | — | — | 1/2% | 6/21% | 3/10% | 3/11% | 5/13% |
| AT5G61380 | TOC1 | 69 kDa | 2/4% | 2/4% | 2/5% | — | 3/7% | — | — | — | 1/2% | 1/2% |
| AT1G53090 | SPA4 | 89 kDa | 2/6% | 5/14% | — | — | — | — | — | — | — | — |
| AT5G18190 | MLK1 | 77 kDa | 2/12% | 2/11% | — | — | — | — | 2/14% | 2/14% | 2/11% | 2/11% |
| AT3G26640 | LWD2 | 39 kDa | 2/21% | 1/21% | — | 1/16% | — | 2/22% | 3/27% | 1/16% | — | — |
| AT1G12910 | LWD1 | 39 kDa | 2/21% | 3/30% | 2/21% | 4/31% | 1/17% | 2/21% | 2/22% | 2/19% | 0/11% | 0/11% |
| AT4G16250 | phyD | 129 kDa | 1/10% | 1/11% | 2/12% | 1/12% | 3/15% | 2/12% | 2/11% | 1/10% | 2/14% | 2/12% |
| AT1G09570 | phyA | 125 kDa | 1/1% | 1/1% | 7/11% | 5/5% | 7/11% | 2/2% | — | 2/2% | 6/7% | 3/4% |
| AT2G25760 | MLK3 | 76 kDa | 1/5% | 1/6% | — | 1/4% | — | 1/4% | 3/13% | 2/9% | 2/9% | 2/9% |
| AT3G03940 | MLK2 | 78 kDa | 1/8% | 1/8% | — | 1/7% | — | 0/2% | 3/20% | 2/13% | 3/13% | 1/9% |
| AT3G46640 | LUX | 35 kDa | 1/3% | 1/3% | 0/10% | 3/19% | 0/10% | 3/23% | 1/3% | 1/3% | 1/3% | 2/11% |
| AT1G17455 | ELF4-L4 | 13 kDa | — | — | — | — | — | — | 1/23% | 1/23% | — | — |
| AT1G72630 | ELF4-L2 | 13 kDa | — | 1/11% | — | 1/13% | — | — | 1/22% | 1/22% | 1/22% | 1/11% |

Proteins co-purified with ELF3 orthologs (AtELF3, SvELF3 and BdELF3, with C-terminal His6-3xFLAG tag) were identified from affinity purification coupled with mass spectrometry (AP-MS) analyses using 12L:12D grown, 10-day-old transgenic plants (in *elf3-2 null* mutant backgrounds) harvested at ZT12.
<sup>a</sup> All listed proteins match 99% protein threshold, minimum number peptides of 2 and peptide threshold as 95%. Proteins not matching the criteria were marked with "—".
<sup>b</sup> Percent coverage for PCH1 is calculated using protein encoded by *At2g16365.2*
<sup>c</sup> These proteins have been noted as frequently identified proteins in AP-MS experiments (see Van Leene, 2015).

We have previously reported proteins that co-precipitated with ELF3 driven from its native promoter using a similar AP-MS methodology (Huang et al., 2016a). When using the 35S promoter driven AtELF3 transgenic line, we were able to generate a curated list of 22 proteins that specifically co-precipitate with AtELF3, including all previously identified proteins, such as all five phytochromes, PHOTOPERIODIC CONTROL OF HYPOCOTYL1 (PCH1) (Huang et al., 2016c), and COP1 (**Table 1**). In addition, we also identified LIGHT-REGULATED WD 2 (LWD2) and SPA1-RELATED 4 (SPA4) as now co-precipitating with AtELF3. These additional interactions may be a result of a combination of altered seedling age, expression level of the ELF3 bait, or tissue-specificity of expression due to these purifications are from tissues where the epitope-tagged transgene is constitutively over-expressed. However, since LWD2 is a known component of the circadian clock (Wu et al., 2008) and SPA4 is a known component of the COP1-SPA complex (Zhu et al., 2008), these interactions are likely to be relevant.

In comparing the list of BdELF3 and SvELF3 co-precipitated proteins with that of AtELF3, we found that neither SvELF3 nor BdELF3 co-precipitated SPA2 and SPA4, components of the COP1-SPA complex. In addition, SvELF3 did not co-precipitate MUT9-LIKE KINASE1, a kinase with roles in chromatin modification and circadian rhythms as AtELF3 did (Huang et al., 2016a; Wang et al., 2015). However, BdELF3 and SvELF3 associated with most of the proteins found in AtELF3 AP-MS (20 out of 22 for BdELF3, 19 out of 22 for SvELF3), in at least one of the replicate purifications from each monocot ortholog AP-MS. Therefore our data suggest that BdELF3 and SvELF3 are integrated into a similar protein-protein interaction network as AtELF3, which likely underlies their ability of broadly complementing *elf3* mutants.

## DISCUSSION

Recent work in diverse plant species has found that the circadian clock plays critical roles in regulating metabolism, growth, photoperiodism, and other agriculturally important traits (Bendix et al., 2015; McClung, 2013; Shor and Green, 2016). While the relevance of the circadian clock to plant fitness is unquestioned, it is unclear if the circadian clock components have conserved function among different plant species. This is particularly true for the majority of clock proteins, whose biological functions are currently poorly understood at the molecular level (Hsu and Harmer, 2014). Also, the divergent modes of growth regulation and photoperiodism between monocots and dicots suggest that the clock evolved to have altered roles in regulating these physiological responses between lineages (Matos et al., 2014; Poire et al., 2010; Song et al., 2014). Here we asked if orthologs of ELF3 from two monocots could complement any of the loss-of-function phenotypes in the model dicot plant *A. thaliana*. In this study we found that ELF3 from either *B. distachyon* or *S. viridis* could complement the hypocotyl elongation, early flowering, and arrhythmic clock phenotype of the *elf3* mutant in *A. thaliana,* despite the variations in protein sequences and evolutionary divergence between monocot and dicot plants. These data suggest that monocot ELF3s can functionally substitute for *A. thaliana* ELF3, albeit with varying efficacy. Since monocot and dicot ELF3 are largely different in the protein sequences, functional conservation of ELF3 orthologs also leads to the next open question of identifying the functional domains within ELF3.

Previously, comparison of ELF3 homologs has identified at least five conserved regions that may be important for function (**Supplemental Figure S1**) (Liu et al., 2001; Saito et al., 2012; Weller et al., 2012). Our multiple sequence alignments also show that at least two regions of AtELF3, namely the N-terminus (AA 1~49) and one middle region (AA 317~389) share many conserved residues with ELF3 orthologs in grasses (**Supplemental Figure S1**). These regions fall within known fragments that are sufficient for binding to phyB (Liu et al., 2001), COP1 (Yu et al., 2008), or ELF4 (Herrero et al., 2012). Consistent with the hypothesis that these conserved regions are critical for proper ELF3 function, a single amino acid substitution (A362V) within this middle region results in defects of ELF3 nuclear localization and changes in the circadian clock period (Anwer et al., 2014). In addition, our protein-protein interaction study and AP-MS analysis show that both monocot ELF3 can form composite ECs (**Figure 5**) and that all three ELF3 homologs interact with an almost identical set of proteins *in vivo* (**Table 1**), further suggesting that one or more of the conserved regions may mediate the binding between ELF3 and its known interacting proteins. Furthermore, the similar pool of ELF3 interacting proteins identified by Bd/SvELF3 AP-MS suggests that the overall conformation of ELF3 ortholog proteins is conserved and that similar complexes and interactions with ELF3 orthologs may form in monocot species. However, whether these interactions form *in planta* and have the same effect on physiology is unclear. For example, *S. viridis* data generated here and public data for *B. distachyon* and *O. sativa*, showed that ELF3 does not cycle under circadian conditions, which differs from Arabidopsis. Further, different from the fact that the clock plays a key role in regulating elongation in *A. thaliana* (Nozue et al., 2007), the circadian clock has no influence on growth in C3 model grass *B. distachyon,* despite robust oscillating expression of putative clock components (Matos et al., 2014). Similarly, ELF3 from rice *(Oryza sativa)* and soybean promotes flowering and senescence (Lu et al., 2017; Saito et al., 2012; Sakuraba et al., 2016; Yang et al., 2013; Zhao et al., 2012), while in *A. thaliana,* ELF3 represses these responses (Liu et al., 2001; Sakuraba et al., 2014; Zagotta et al., 1992), which suggests significant rewiring of ELF3 regulated photoperiodic responses of flowering between short-day (rice/soybean) and long-day (*A. thaliana*) plants. Alternatively, ELF3 may form distinct interactions and complexes in monocot species that were not identified in our trans-species complementation analysis. Clearly, further work is required to understand ELF3 function in monocots beyond the studies presented here.

In addition to the molecular characterization of ELF3, our analysis of circadian-regulated genes in *S. viridis* after photo- and thermo-entrainment found significant differences in the behavior of the clock when compared to other monocots. Although the number of circadian regulated genes is comparable to studies done in corn and rice after photo-entrainment (between 10-12%) (Filichkin et al., 2011; Khan et al., 2010), we found that very few genes (~1%) continue to cycle after release from temperature entrainment in *S. viridis* (**Supplemental Figure S4**) when compared to rice (~11%) (Filichkin et al., 2011). This may reflect a fundamental difference in how the clock interfaces with temperature between these monocot species. Furthermore, proportions of circadian regulated genes upon photo-entrainment in all three monocot plants (**Supplement Figure S4**) (Filichkin et al., 2011; Khan et al., 2010) are much smaller than the approximately 30% reported for *A. thaliana* (Covington et al., 2008), suggesting the divergence of clock functions through evolution or domestication. Further comparisons of circadian responses among monocots or between monocots and dicots will help to determine the molecular underpinning of these differences.

In summary, we find that BdELF3 and SvELF3 form similar protein complexes *in vivo* as AtELF3, which likely allows for functional complementation of loss-of-function of *elf3* despite relatively low sequence conservation. We also present an online query tool, Diel Explorer that allows for exploration of circadian gene expression in *S. viridis*, which illustrate fundamental differences in clock function among monocots and between monocots and dicots. Collectively, this work is a first step toward functional understanding of the circadian clock in two model monocots, *S. viridis* and *B. distachyon.*

## MATERIALS AND METHODS

### Plant materials and growth conditions

For *A. thaliana,* wild type (Columbia-0) and *elf3-2* plants carrying the CCA1::LUC reporter were described previously (Nusinow et al., 2011; Pruneda-Paz et al., 2009). Seeds were surface sterilized and plated on 1/2x Murashige and Skoog (MS) basal salt medium with 0.8% agar + 1% (w/v) sucrose. After 3 days of stratification, plates were placed horizontally in a Percival incubator (Percival Scientific, Perry, IA), supplied with 80 μmol m^-2^ sec^-1^ white light and set to a constant temperature of 22°C. Plants were grown under 12 h light / 12 h dark cycles (12L:12D) for 4 days (for physiological experiments) or for 10 days (for AP-MS) before assays.

For *S. viridis* circadian expression profiling by RNA-seq, seeds were stratified for 5 days at 4°C before being moved to entrainment conditions. Plants were grown under either LDHH or LLHC (L: light, D: dark, H: hot, C: cold) entrainment condition, and then sampled for RNA-seq in constant light and constant temperature (32°C) conditions (F, for free-running) every 2 hours for 48 hours. Light intensity was set to 400 μmol m^-2^ sec^-1^ white light. In LDHH-F, stratified *S. viridis* seeds were grown for 10 days under 12L:12D and constant temperature (32°C) before sampling in constant light and constant temperature. In LLHC-F, stratified *S. viridis* seeds were grown for 10 days under constant light conditions and cycling temperature conditions 12 h at 32°C (subjective day) / 12 h at 22°C (subjective night) before sampling in constant conditions. Two experimental replicates were collected for each entrainment condition.

### Setaria circadian RNA-seq

The second leaf from the top of seventeen *S. viridis* plants was selected for RNA-seq sampling at each time point for each sampling condition. Five replicate samples were pooled after being ground in liquid nitrogen and resuspended in lithium chloride lysis binding buffer (Wang et al., 2011a). RNA-seq libraries from leaf samples were constructed according to the previous literature (Wang et al., 2011a) with one major modification. Rather than extracting RNA then mRNA from ground leaf samples (Wang et al., 2011a), mRNA was extracted directly from frozen ground leaf samples similar to the method described in (Kumar et al., 2012), except that two additional rounds of wash, binding, and elution steps after treatment with EDTA were necessary to remove rRNA from samples. mRNA quantity was assessed using a Qubit with a Qubit RNA HS Kit and mRNA quality was assessed using a Bioanalyzer and Plant RNA PiCO chip. 96 library samples were multiplexed 12 per lane, for a total of 8 lanes of Illumina Hiseq 2000 sequencing. Paired end 101 bp Sequencing was done at MOgene (St. Louis, MO). Raw data and processed data can be found on NCBI’s Gene Expression Omnibus (GEO; (Barrett et al., 2013; Edgar et al., 2002)) and are accessible with identification number GSE97739 (https://www.ncbi.nlm.nih.gov/geo/query/acc.cgi?acc=GSE97739).

RNA-seq data was trimmed with BBTools (v36.20) using parameters: ktrim=r k=23 mink=11 hdist=1 tpe tbo ktrim=l k=23 mink=11 hdist=1 tpe tbo qtrim=rl trimq=20 minlen=20 (Bushnell, 2016). Any parameters not specified were run as default. Before trimming we had 1,814,939,650 reads with a mean of 18,905,621 reads per sample and a standard deviation of 2,875,187. After trimming, we have 1,646,019,593 reads with a mean of 17,146,037 reads per sample and a standard deviation of 2,411,061. Kallisto (v 0.42.4; (Bray et al., 2016)) was used to index the transcripts with the default parameters and the *S. viridis* transcripts fasta file (Sviridis_311_v1.1) from Phytozome (Goodstein et al., 2012). The reads were quantified with parameters:-t 40 -b 100. Any parameters not specified were run as default. Kallisto output was formatted for compatibility with JTKCycle (v3.1; (Hughes et al., 2010)) and circadian cycles were detected. To query the *S. viridis* expression data we developed Diel Explorer. The tool can be found at http://shiny.bioinformatics.danforthcenter.org/diel-explorer/. Underlying code for Diel Explorer is available on Github (https://github.com/danforthcenter/diel-explorer).

### Plasmid constructs and generation of transgenic plants

Coding sequences (without the stop codon) of *AtELF3* (*AT2G25930*) and *SvELF3a* (*Sevir.5G206400.1*) were cloned into the pENTR/D-TOPO vector (ThermoFisher Scientific, Waltham, MA), verified by sequencing and were recombined into the pB7HFC vector (Huang et al., 2016a) using LR Clonease (ThermoFisher Scientific, Waltham, MA). Coding sequence of BdELF3 (*Bradi2g14290.1*) was submitted to the U.S. Department of Energy Joint Genome Institute (DOE-JGI), synthesized by the DNA Synthesis Science group, and cloned into the pENTR/D-TOPO vector. Sequence validated clones were then recombined into pB7HFC as described above. The pB7HFC-At/Bd/SvELF3 constructs were then transformed into *elf3-2* [*CCA1::LUC*] plants by the floral dip method (Zhang et al., 2006). Homozygous transgenic plants were validated by testing luciferase bioluminescence, drug resistance, and by PCR-based genotyping. All primers used in this paper were listed in **Supplemental Table S1**.

### Hypocotyl and flowering time measurement

20 seedlings of each genotype were arrayed and photographed with a ruler for measuring hypocotyl length using the ImageJ software (NIH) (Schneider et al., 2012). The procedure was repeated three times. For measuring flowering time, 12 plants of each genotype were placed in a random order and were grown under the long day condition (light : dark = 16 : 8 hours). The seedlings were then observed every day at 12:00 PM; the date on which each seedling began flowering, indicated by the growth of a ~1 cm inflorescence stem, was recorded along with the number of rosette leaves produced up to that date. ANOVA analysis with Bonferroni correction was measured using GraphPad Prism version 6.00 (GraphPad Software, La Jolla California USA, www.graphpad.com).

### Circadian assays in *A. thaliana*

Seedlings were transferred to fresh 1/2x MS plates after 5 days of entrainment under the 12L:12D condition and sprayed with sterile 5 mM luciferin (Gold Biotechnology, St. Louis, MO) prepared in 0.1% (v/v) Triton X-100 solution. Sprayed seedlings were then imaged in constant light (70 μmol m^-2^ sec^-1^, wavelengths 400, 430, 450, 530, 630, and 660 set at intensity 350 (Heliospectra LED lights, Göteborg, Sweden)). Bioluminescence was recorded after a 120-180s delay to diminish delayed fluorescence (Gould et al., 2009) over 5 days using an ultra-cooled CCD camera (Pixis 1024B, Princeton Instruments) driven by Micro-Manager software (Edelstein et al., 2010; Edelstein et al., 2014). The images were processed in stacks by Metamorph software (Molecular Devices, Sunnyvale, CA), and rhythms determined by fast Fourier transformed non-linear least squares (FFT-NLLS) (Plautz et al., 1997) after background subtraction using the interface provided by the Biological Rhythms Analysis Software System 3.0 (BRASS) available at http://www.amillar.org.

### Yeast two-hybrid analysis

Yeast two-hybrid assays were carried out as previously described (Huang et al., 2016a). In brief, the DNA binding domain (DBD) or activating domain (AD)-fused constructs were transformed using the Li-Ac transformation protocol (Clontech) into *Saccharomyces cerevisiae* strain Y187 (MATα) and the AH109 (MATa), respectively. Two strains of yeast were then mated to generate diploid with both DBD and AD constructs. Protein-protein interaction was tested in diploid yeast by replica plating on CSM –Leu –Trp –His media supplemented with extra Adenine (30mg/L final concentration) and 2mM 3-Amino-1,2,4-triazole (3AT). Pictures were taken after 4-day incubation at 30 °C. All primers used for cloning plasmid constructs were listed in **Supplemental Table S1**.

### Protein extraction and western blotting

Protein extracts were made from 10-day-old seedlings as previously described (Huang et al., 2016a) and loaded 50 μg to run 10% SDS-PAGE. For western blots, all of the following primary and secondary antibodies were diluted into PBS + 0.1% Tween and incubated at room temperature for 1 hour: anti-FLAG®M2-HRP (Sigma, A8592, diluted at 1:10,000) and anti-Rpt5-rabbit (ENZO Life Science, BML-PW8245-0025, diluted at 1:5000) and anti-Rabbit-HRP secondary antibodies (Sigma, A0545, diluted at 1:10,000).

### Affinity purification and mass spectrometry

Protein extraction methods and protocols for AP-MS were described previously (Huang et al., 2016a; Huang et al., 2016b; Huang and Nusinow, 2016a; Huang et al., 2016c)}. In brief, transgenic seedlings carrying the At/Bd/SvELF3-HFC constructs were grown under 12L:12D conditions for 10 days and were harvested at dusk (ZT12). 5 grams of seedlings were needed per replicate to make protein extracts, which underwent tandem affinity purification utilizing the FLAG and His epitopes of the fusion protein. Purified samples were reduced, alkylated and digested by trypsin. The tryptic peptides were then injected to an LTQ-Orbitrap Velos Pro (ThermoFisher Scientific, Waltham, MA) coupled with a U3000 RSLCnano HPLC (ThermoFisher Scientific, Waltham, MA) with settings described previously (Huang et al., 2016a).

### AP-MS data analysis

Data analysis was done as previously described (Huang et al., 2016a). The databases searched were TAIR10 database (20101214, 35,386 entries) and the cRAP database (http://www.thegpm.org/cRAP/). Peptide identifications were accepted if they could be established at greater than 95.0% probability and the Scaffold Local FDR was <1%. Protein identifications were accepted if they could be established at greater than 99.0% probability as assigned by the Protein Prophet algorithm (Keller et al., 2002; Nesvizhskii et al., 2003). A full list of all identified proteins (reporting total/exclusive unique peptide count and percent coverage) can be found in **Supplemental Table S2**. The mass spectrometry proteomics data have been deposited to the ProteomeXchange Consortium (Vizcaino et al., 2014) via the PRIDE partner repository with the dataset identifier PXD006352 and 10.6019/PXD006352.

## Supplemental Data

**Supplemental Table S1.** List of all primers used.

**Supplemental Table S2.** A full list of At/Bd/SvELF3 associated proteins identified from AP-MS.

**Supplemental Figure S1.** Multiple sequence alignments of ELF3 orthologs

**Supplemental Figure S2.** Diel and circadian expression of *BdELF3* from the DIURNAL database.

**Supplemental Figure S3.** Example of the Diel Explorer interface.

**Supplemental Figure S4.** Summary of circadian regulated genes in *S. viridis.*

**Supplemental Figure S5.** Circadian expression of selected *A. thaliana* clock genes from the DIURNAL database.

**Supplemental Figure S6.** Anti-FLAG western of ELF3 transgenic lines used for complementation analysis.

**Supplemental Figure S7.** Relative Amplitude Error vs period plots of transgenic *elf3-2* plants expressing At/Bd/SvELF3.

